# Cell Ploidy Modulates Nucleolar Size via Dosage of Nucleolar Organizing Regions

**DOI:** 10.64898/2026.02.13.705825

**Authors:** Tianhao Bi, Varuni Rastogi, Maged Zeineldin, Blake A. Johnson, Yi Dong, Mulin Li, Tatianna C. Larman

## Abstract

Abnormal nucleolar morphology is a pathognomonic histologic feature of cancer cells. Nucleoli are biomolecular condensates and cytogenetic entities that form around nucleolar organizing regions (NORs), located on the short arms of acrocentric chromosomes. Given that aneuploidy is pervasive in human cancers and can involve NOR-bearing chromosomes, we investigated how aneuploidy-driven changes in NOR dosage impact nucleolar morphology. We used reversine, a spindle assembly checkpoint inhibitor, to induce chromosomal instability and complex aneuploidy in DLD1 colorectal cancer cells (near-euploid) and normal human colon organoids (colonoids, euploid). High resolution 3D imaging and quantitative analyses of reversine-treated cells showed overall doubling of total nucleolar volume per nucleus. Reversine-associated nucleolar enlargement occurred independently of global protein synthesis, and thus was not solely driven by increases in biosynthetic demand. The volume of individual nucleoli showed a direct linear correlation with the number of associated NORs, as revealed by 3D co-localization with NOR-specific FISH probes. These findings establish NOR dosage as a determinant of nucleolar morphology and metric of genomic instability, providing a new paradigm by which aneuploidy can modulate spatial chromosomal constraints and nuclear architecture in cancer.

**Significance statement:** Cancer cells often show enlarged and abnormal nucleoli. We show that aneuploidy, a common feature of cancer, can directly reshape nucleolar morphology by altering the copy number of acrocentric chromosomal regions around which nucleoli form, NORs. Using high-resolution 3D imaging in cell lines and patient-derived colonoids, we find that nucleolar number and volume directly correlate with NOR number, independent of overall protein synthesis. Individual nucleolar volume scales with the number of 3D NOR associations, and prominent single nucleoli arise when NORs accumulate. These results provide evidence that aneuploidy can lead to aberrant nucleolar morphologies seen in many human cancers.

## Introduction

Aberrant nucleolar morphologies, particularly enlarged and prominent single nucleoli, are among the key cytologic features of malignancy that pathologists rely on diagnostically (1) (2) (3). In fact, the histologic grading of some tumor types is solely based upon nucleolar features, which correlate directly with patient prognosis (4). Nucleoli are dynamic and membraneless nuclear domains that play a central role in ribosomal RNA (rRNA) synthesis, ribosome biogenesis, and regulation of cellular homeostasis (5) (6). Importantly, nucleoli are also cytogenetic entities and form around nucleolar organizing regions (NORs), which are located on the short arms of the five human acrocentric chromosomes (chromosomes 13, 14, 15, 21 and 22) and comprised of tandem arrays of ribosomal DNA (rDNA) (7).

More than a century since their first description, the molecular basis for prominent nucleoli in cancer is not fully understood (8) (9). Nucleolar enlargement in cancer and inflammation has been descriptively attributed to increased ribosomal biogenesis needed for cell proliferation, and has also been associated with stemness and oncogene activation (10) (11) (12) (13). However, there is mounting evidence that nucleoli have other underexplored cellular roles, including in cell-cycle regulation, stress response, genome stability, proteostasis, and in modulation of gene expression via protein sequestration (14) (15) (16) (17).

Aneuploidy, defined as the gain or loss of whole chromosomes, characterizes more than 90% of human solid tumors (18), and may conceivably involve acrocentric chromosomes and NORs. However, owing to their highly repetitive nature, we have lacked consensus normal human reference sequences for the NOR-bearing short arms of acrocentric chromosomes (7) (19). This lack of a reference sequence has limited our ability to bioinformatically mine existing cancer genomic datasets with abnormal nucleolar morphologies for rDNA quantification and/or characterization of NOR-associated copy number. Consequently, whether NOR dosage correlates with nucleolar phenotypes in human cancer has not been investigated to date.

In this study, we investigate whether modulation of NOR copy number, via aneuploidy involving NOR-bearing chromosomes, impacts nucleolar morphology in human cells. Using monopolar spindle 1 (MPS1) inhibitor reversine to induce chromosomal instability and complex aneuploidy in near-euploid colorectal cancer cells and normal human colon epithelial organoids, we then employed high-resolution 3D confocal imaging and surface rendering to quantify nucleolar number and volume at the single-cell level. Nucleolar size scaled proportionally with total genomic content and cellular volume in reversine-treated cells, reflecting a regulated structural response to complex aneuploidy. This nucleolar enlargement occurred independently of protein synthesis levels, suggesting that biosynthetic capacity is not the sole driver of nucleolar size. This increased nucleolar volume directly correlated with direct NOR associations, evidence that aneuploidy-associated modulation of NOR dosage is a key determinant of nucleolar phenotypes.

These results reveal a previously unrecognized mechanism linking aneuploidy to nucleolar changes common in cancer. By establishing NOR amplification as a genomic determinant of nucleolar enlargement in human cells, this work provides new insights into a non-genomic consequence of chromosomal instability and aneuploidy on nucleolar architecture.

## Results

### Reversine treatment increases karyotype complexity in cancer cells and normal human colonoids

To examine how altered chromosome number affects nucleolar morphology, we perturbed chromosome segregation in two epithelial *in vitro* lines with near-normal karyotypes: DLD1 colorectal cancer cells, and patient-derived normal colon organoids (colonoids). Our goal was to pharmacologically induce near-tetraploidy in order to alter dosage of the five human acrocentric chromosomes which harbor NORs (20). Indeed, in a previous study using normal colonoids line treated with MPS1 inhibitor reversine, 47% (47 of 101 aneuploid single cell karyotypes) harbored aneuploidy involving at least one acrocentric chromosome (21). Thus, both DLD1 cells and colonoids were exposed to reversine for 24 hours, washed, recovered in drug-free medium for another 24 hours, and analyzed by metaphase chromosome spreads and high-resolution 3D confocal imaging for nucleolar morphology (Figure 1*A*).

**Figure 1.**
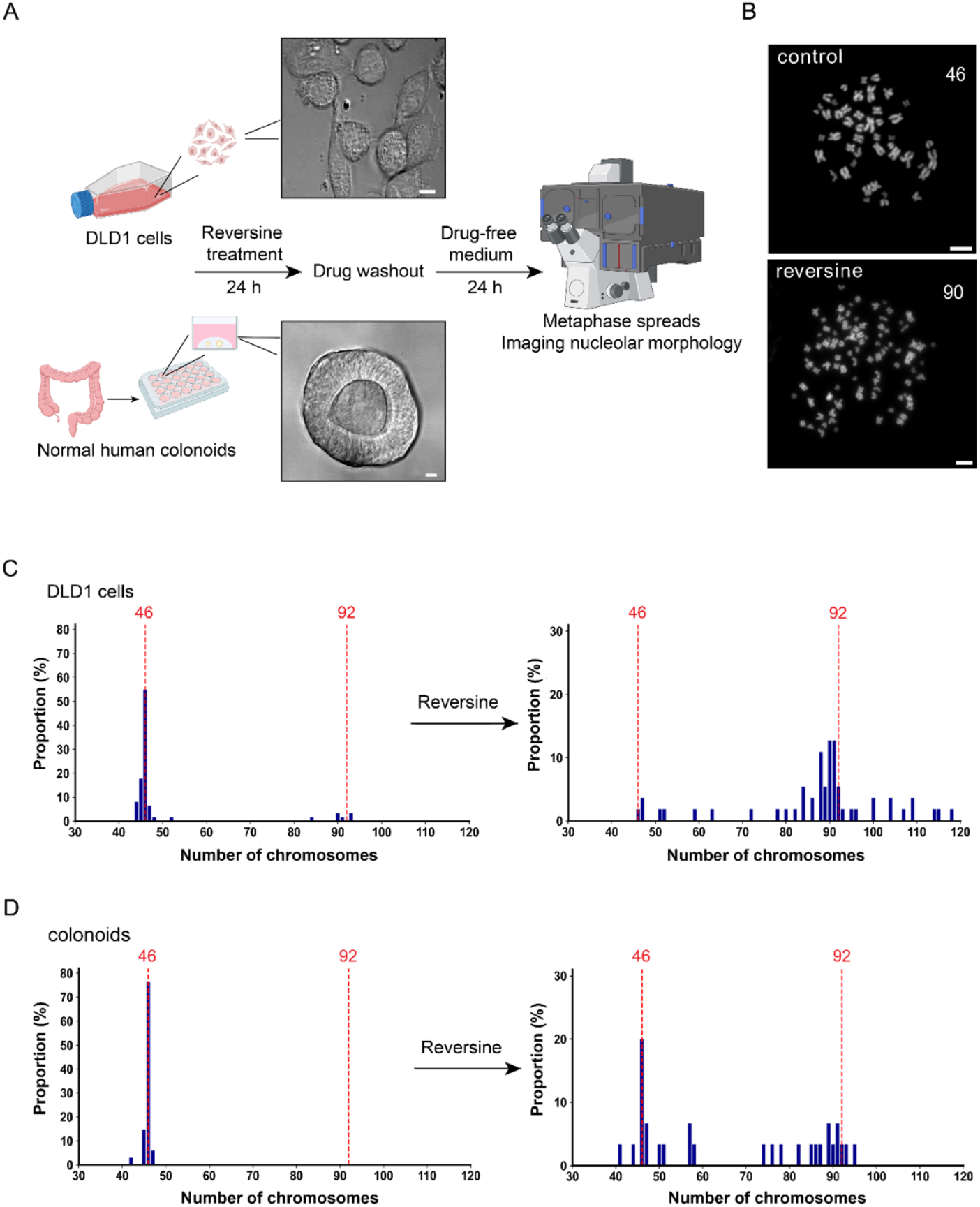
Reversine induces complex aneuploidy. (**A**) Experimental workflow and representative images of DLD1 cells and human colonoids. DLD1 cells and colonoids were treated with MPS1 inhibitor reversine, followed by metaphase spread chromosome analysis and 3D imaging. Scale bars, 10 µm. Image was created using BioRender. (**B**) Representative images of metaphase spreads from vehicle-control (top) and reversine-treated (bottom) DLD1 cells. 46 and 90 represent the number of chromosomes in each spread. Scale bars, 10 µm. (**C**) Karyotype distributions of vehicle-control DLD1 cells (left) and reversine-treat DLD1 cells (right). (n=62 for control DLD1 cells, n=55 for reversine-treated DLD1 cells). (**D**) Karyotype distributions of vehicle-control colonoids (left) and reversine-treated colonoids (right). (n=34 for control colonoids, n=30 for reversine-treated colonoids).

Metaphase spreads showed markedly increased chromosomal complexity after reversine treatment, as expected (Figure 1*B*). In DLD1 cells, chromosome counts shifted from predominantly 46 in controls to an increased range (up to 118 chromosomes) after reversine treatment, with 67% of spreads showing near-tetraploid karyotypes (in the range of 82-102 chromosomes) (Figure 1*C*). In colonoids, chromosome counts similarly shifted from mostly 46 in controls to up to 95 after reversine treatment, and 40% of spreads fell within the near-tetraploid range (Figure 1*D*). Moreover, live cell imaging on DLD1 reporter cell line stably-expressed mCherry-CAAX (to delineate the plasma membrane), H2B-mNeon (to mark the nucleus) and BFP-NPM1 (to label nucleoli) (22) (23) also confirmed ongoing chromosomal instability and mitotic errors in the reversine-treated group. With reversine treatment, more than 70% of mitotic errors were monopolar (Supplementary Figure 1*A-D*). These results confirm that reversine treatment induces complex aneuploidy and frequent near-tetraploid karyotypes in both normal and cancer epithelial lines.

### Reversine-treatment results in overall doubling of total nucleolar volume

To determine how these chromosomal alterations affect nucleolar morphologies, we performed high-resolution 3D confocal imaging on DLD1 cells and colonoids. Cells were stained with Hoechst to label DNA and identify nuclear boundaries, and with Nucleolar ID® (binds to ribosomal RNA) to selectively mark nucleoli (24). Z-stack images were acquired and subsequently reconstructed in Imaris software for volumetric segmentation and analysis (Figure 2*A*). Specifically, we extracted key structural parameters including nuclear volume, the number of nucleoli per nucleus, the volume of individual nucleoli, and total nucleolar volume per nucleus. Both control and reversine-treated DLD1 cells exhibited a normally distributed range of nucleolar numbers per nucleus; however, reversine-treated cells showed a clear rightward shift, with a significant increase in mean nucleolar count (from 2.9 to 4.5) and near doubling of the range of nucleolar number per nucleus (from 1-6 to 1-10) (Figure 2*B*).

**Figure 2.**
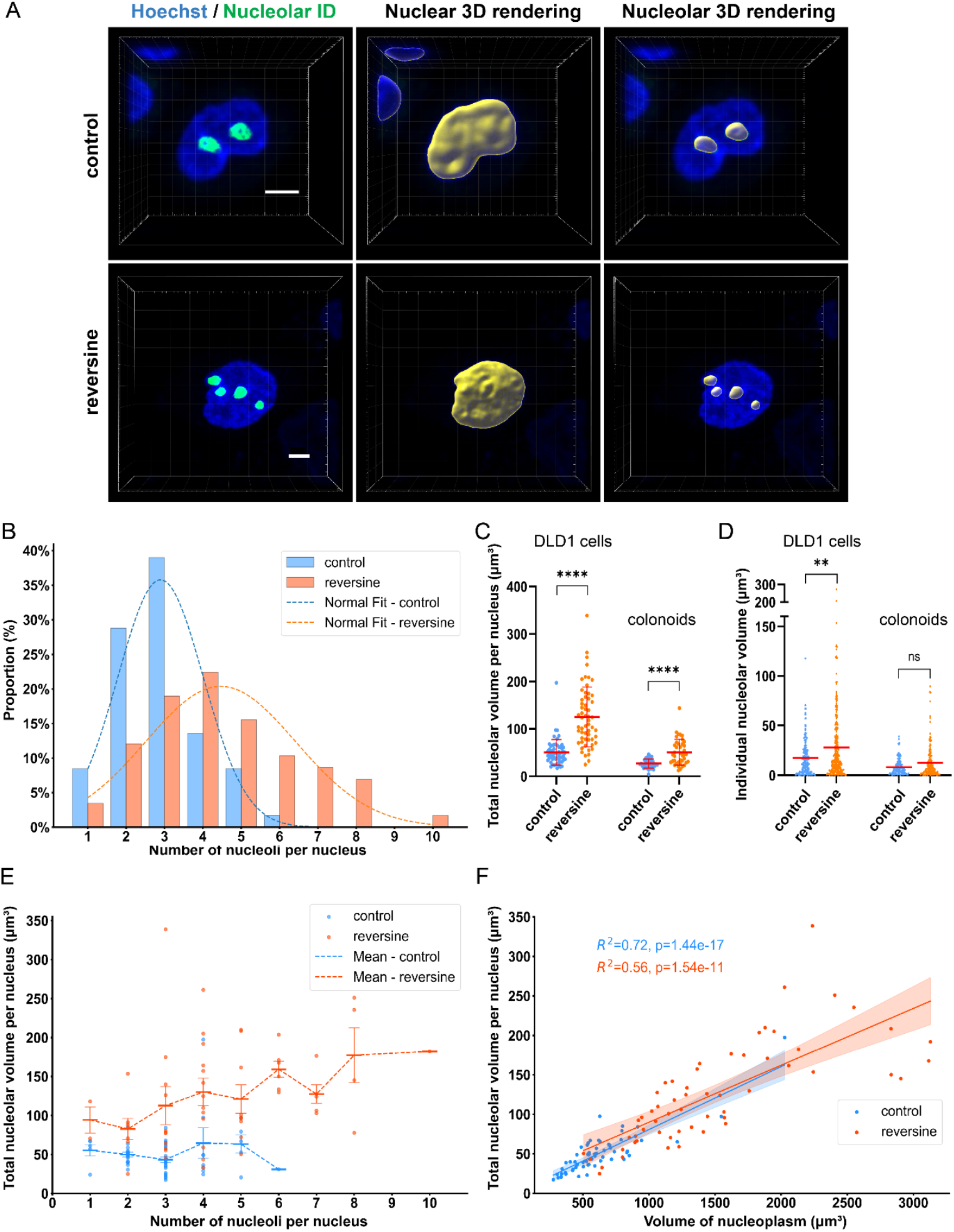
Reversine treatment results in overall doubling of nucleolar number and volume. (**A**) Representative 3D confocal images showing Hoechst-labeled nuclei and Nucleolar ID®-stained nucleoli in vehicle-control (upper) and reversine-treated (bottom) DLD1 cells.Volumetric surface renderings were quantified using Imaris software. Scale bars, 5 µm. (**B**) Histogram showing the distribution of number of nucleoli per nucleus in control and reversine-treated DLD1 cells. Dashed lines represent the normal distribution fits based on mean and standard deviation. (**C**) Total nucleolar volume per nucleus in DLD1 cells (left) and colonoids (right) between control and reversine-treated groups. Lines indicate the mean, and error bars represent the SD. **** p < 0.0001 (Mann-Whitney test). (n=59 for control DLD1 cells, n=58 for reversine-treated DLD1 cells; n=40 for both control and reversine-treated colonoids). (**D**) Individual nucleolar volume in DLD1 cells (left) and colonoids (right) between control and reversine-treated groups. Lines indicate the mean. **p < 0.01, ns means not significant (Mann-Whitney test). (n=171 for control DLD1 cells, n=259 for reversine-treated DLD1 cells; n=133 for control colonoids, n=162 for reversine-treated colonoids). (**E**) Total nucleolar volume per nucleus plotted against nucleolar number per nucleus in DLD1 cells. Each point represents one nucleus. Lines indicate the mean, and error bars represent the SEM. (**F**) Linear correlation analysis showing total nucleolar volume per nucleus and nucleoplasm volume in DLD1 cells. Each point represents an individual nucleus; lines represent linear regression; shaded areas indicate 95% confidence intervals.

Not only did the number of individual nucleoli increase, but the total nucleolar volume per nucleus increased compared to control DLD1 cells. Individual nucleolar volume was also significantly larger in the reversine-treated group compared to controls (Figure 2*C-D*, left). However, within each group, including both control and reversine-treated, cells with varying numbers of nucleoli maintained relatively consistent total nucleolar volume (Figure 2*E*). This suggests that when the karyotype is stable, the total nucleolar volume within a nucleus is independent of nucleolar number.

Nucleoplasm volume, quantified by subtracting nucleolar volume from total nuclear volume, was also increased in reversine-treated cells (Supplementary Figure 2*A*, left). Moreover, total nucleolar volume correlated linearly with nucleoplasm volume (Figure 2*F*). The overall total nucleolar to nucleoplasm volume ratios were conserved between control diploid and reversine-treated near-tetraploid states (Supplementary Figure 2*B*, left), suggesting that nucleolar expansion scales proportionally with genomic content.

We next used colonoids to assess whether nucleolar-ploidy scaling also occurs in normal epithelium. We observed similar results, with nucleolar number and total nucleolar volume increasing after reversine treatment (Figure 2*C* right, Supplementary Figure 2*C-D*). We also measured an increase in nucleoplasm volume; moreover, total nucleolar volume scaled linearly with nucleoplasm volume, and the ratio of nucleolar to nucleoplasm volume remained unchanged after reversine treatment. These data indicate that nucleolar growth after reversine treatment scales proportionally with surrounding nucleoplasm even in non-neoplastic epithelial cells (Supplementary Figure 2*A-B* right, Supplementary Figure 2*E*). This is consistent with prior observations that nuclear-to-nucleolar volume ratios are not quantitatively different between normal tissue, hyperplastic tissue, benign tumors, or malignant tumors (25). Although the average volume of individual nucleoli did not show statistically significant differences between control and reversine-treated colonoids, we did observe emergence of abnormally large individual nucleoli in the reversine-treated group (Figure 2*D*, right).

Together, these results demonstrate that from diploid to near-tetraploid states in both normal and malignant epithelial cells, both the number and the volume of nucleoli increase while volumetric scaling is preserved.

### Nucleolar enlargement in reversine treated cells is not associated with increased protein synthesis

Nucleolar size is impacted by cellular biosynthetic activity, particularly rRNA transcription and protein synthesis, and it is accepted that enlarged nucleoli reflect rapidly proliferating and/or metabolically active cells (10). We next quantified global protein synthesis of individual cells to measure the contribution, if any, of increased translation to the nucleolar enlargement phenotypes 24 hours after reversine induction of complex aneuploidy.

We used the O-propargyl-puromycin (OPP) incorporation assay, which labels nascent polypeptide chains, as a proxy for translational activity (26). To simultaneously visualize cellular compartments including the cell membrane, nucleus, and nucleoli, we generated a stable DLD1 reporter cell line via lentiviral transduction expressing mCherry-CAAX, H2B-mNeon and BFP-NPM1. To verify the specificity of OPP labeling for nascent protein synthesis, cycloheximide was used as a negative control, which blocks translation elongation (Figure 3*A*). This approach enabled high-resolution 3D quantification of nucleolar morphology and direct correlation with translation output in the same cells.

**Figure 3.**
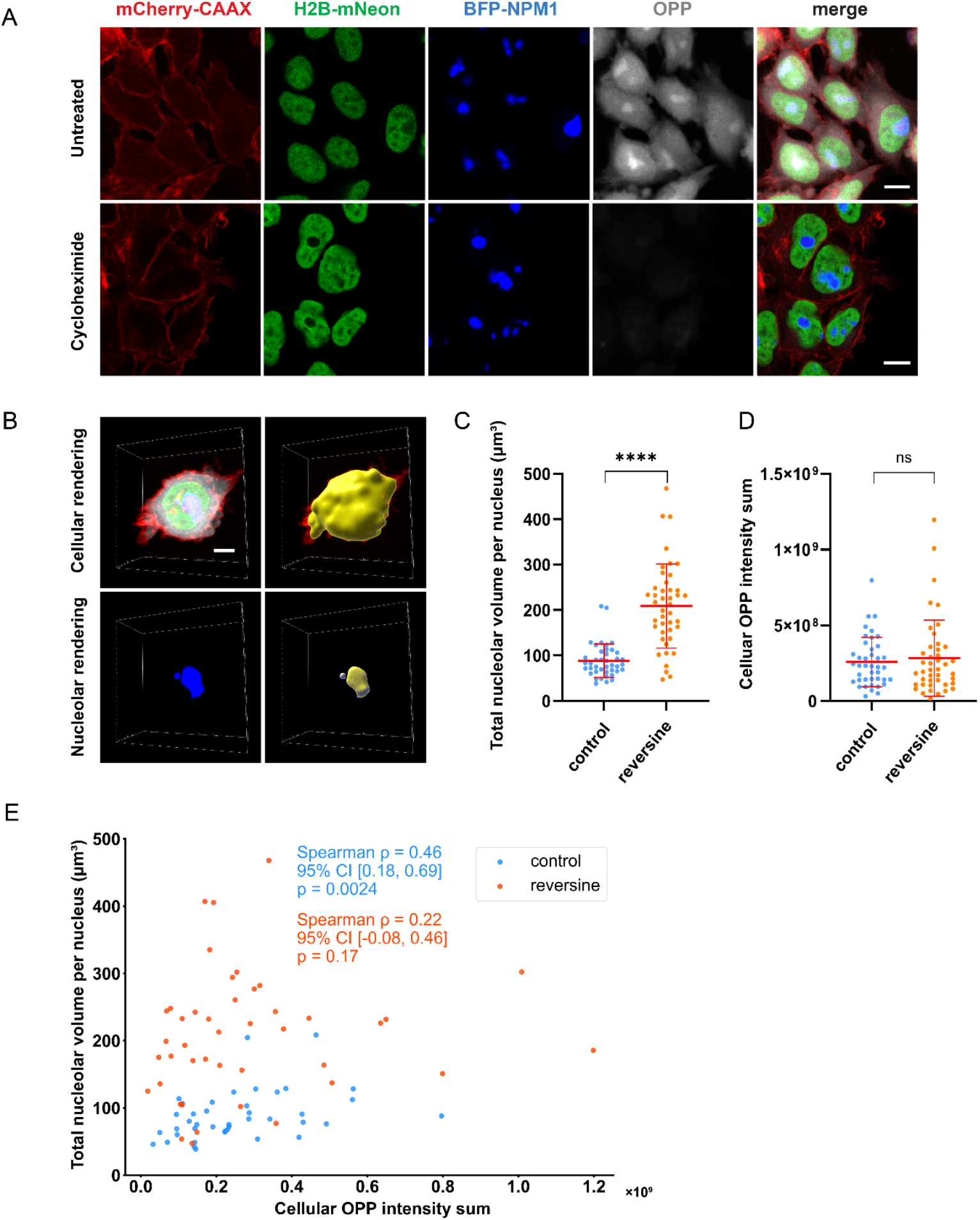
Nucleolar enlargement in reversine-treated cells is not associated with increased global protein synthesis. (**A-B**) Representative images of DLD1 cells stably expressing mCherry-CAAX, H2B-mNeon, and BFP-NPM1 after O-propargyl-puromycin (OPP) labeling. (**A**) Representative confocal images showing OPP-labelled DLD1 cells, with and without cycloheximide treatment (negative control). Scale bars, 10 µm. (**B**) Volumetric surface renderings created using Imaris. Scale bars, 5 µm. (**C-D**) Quantification of total nucleolar volume per nucleus (**C**) and global protein synthesis measured by OPP total cellular fluorescence intensity (**D**) in control versus reversine-treated DLD1 cells. Lines indicate the mean, and error bars represent the SD. ****p < 0.0001, ns means not significant (Mann-Whitney test). (n=41 for control DLD1 cells, n=43 for reversine-treated DLD1 cells). (**E**) Scatter plots showing the relationship between cellular OPP intensity sum and total nucleolar volume per nucleus. Each point represents an individual cell. Spearman rank correlation coefficients (ρ), two-sided p values, and 95% bootstrap confidence intervals are indicated.

Three-color, high-resolution 3D confocal imaging was performed on DLD1 cells status post OPP incorporation, and nucleolar volumes were quantified using segmentation methods (Imaris software) (Figure 3*B*). We observed a significant increase in nucleolar volume per nucleus following reversine treatment, as expected (Figure 3*C*). However, quantification of cellular OPP fluorescence intensity sum did not show a significant difference in global protein synthesis between control and reversine-treated cells (Figure 3*D*). Spearman rank correlation analysis revealed a positive association between cellular OPP intensity sum and total nucleolar volume in control cells (ρ = 0.46, p = 0.0024; 95% CI [0.18, 0.69]), indicating that increased translational activity is associated with variations in nucleolar volume under basal conditions, consistent with previous studies showing that nucleolar size directly reflects the metabolic activity of the cell (27). In contrast, no statistically significant association was detected between cellular OPP intensity sum and nucleolar volume in reversine-treated cells, as the confidence interval spanned zero (ρ = 0.22, p = 0.17; 95% CI [−0.08, 0.46]) (Figure 3*E*). When OPP fluorescence intensity was normalized to cellular volume, OPP intensity was even lower in reversine-treated cells compared to control cells (Supplementary Figure 3*A*). There was no significant correlation between cellular volume–normalized OPP intensity and total nucleolar volume per nucleus in either control cells (ρ = 0.12, p = 0.47; 95% CI [−0.21, 0.42]) or reversine-treated cells (ρ = −0.03, p = 0.86; 95% CI [−0.31, 0.26]) (Supplementary Figure 3*B*). These results suggest that while global metabolic activity is correlated with nucleolar size under basal conditions, the increased spectrum of nucleolar enlargement following reversine treatment is not associated with increased translational activity.

### Nucleolar volume directly correlates with number of 3D NOR associations

To directly test whether nucleolar enlargement in near-tetraploid cells results from increased nucleolar organizing region (NOR) dosage, we performed 3D ImmunoFISH in DLD1 cells. We combined immunofluorescent staining of nucleolin, a nucleolar phosphoprotein that marks nucleolar structure, with DNA fluorescence *in situ* hybridization (FISH) targeting NOR loci. Specifically, we used a probe targeting the distal junction (DJ) NOR sequences, located immediately downstream of rDNA arrays (28, 29). These DJ sequences are highly conserved across all five human acrocentric chromosomes, allowing robust detection of NOR-associated sites regardless of chromosomal origin (29) (29, 30). Of note, the number of discrete NOR foci per nucleus did not vary appreciably across cell cycle stages under our imaging conditions (Supplementary Figure 4*A-B*). Therefore, we used the count of NOR foci to represent NOR dosage at the single-nucleus level.

High-resolution 3D confocal imaging was performed following staining at the same timepoint as our live imaging experiment, and volumetric analysis was carried out using the segmentation strategy employed in prior experiments (Figure 4*A*; Supplemental Videos 1 and 2). This approach enabled us to quantify both nucleolar volume and NOR copy number per nucleus, as well as the number of NORs associated with individual nucleoli.

**Figure 4.**
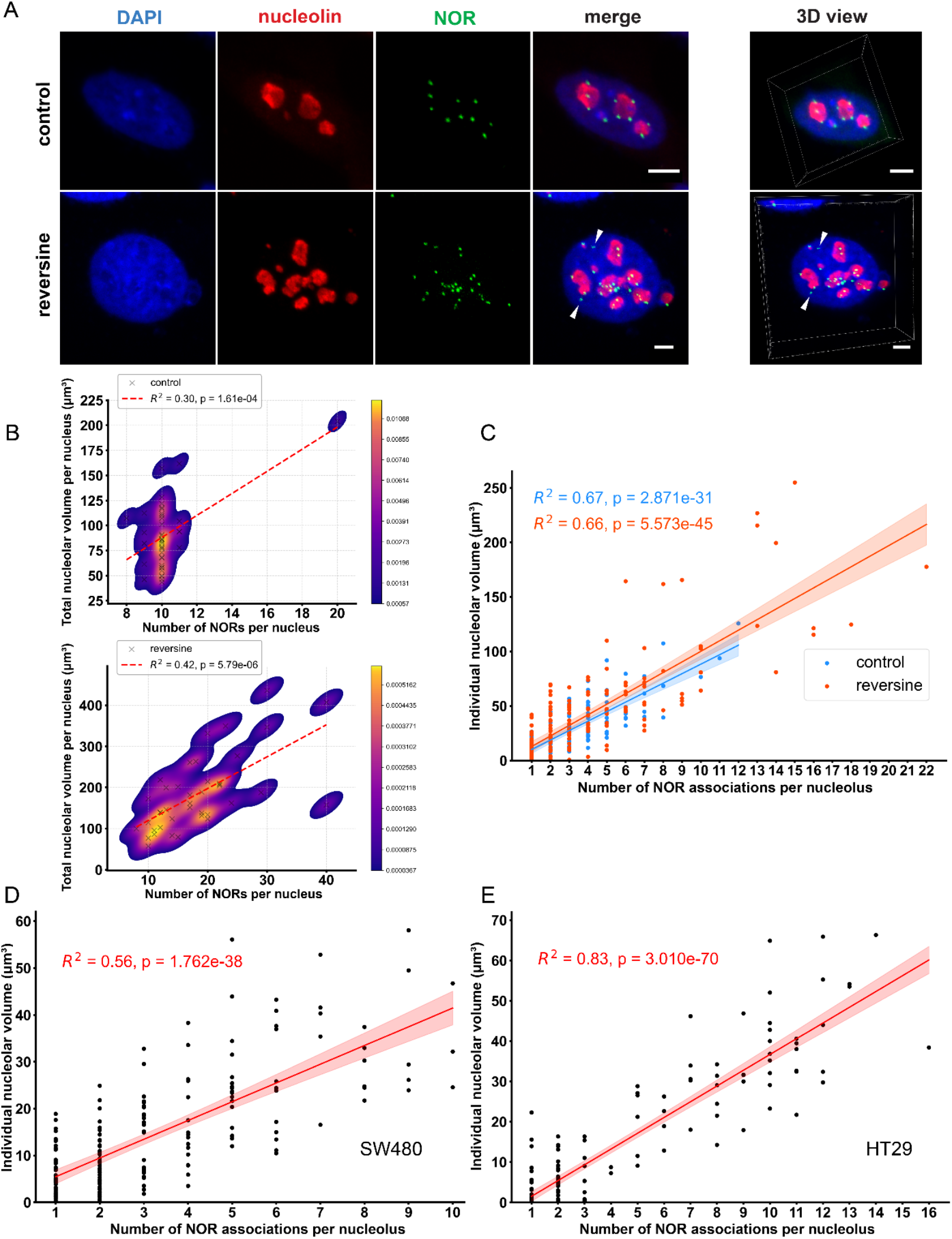
NOR number modulates nucleolar volume. (**A**) Representative 3D ImmunoFISH images of DLD1 cells labeled with anti-nucleolin antibody (red, nucleoli) and distal junction (DJ)-specific FISH probes targeting NOR loci (green). DNA stained with DAPI (blue). White arrowheads denote non-nucleolar associated NORs. Scale bars, 5 µm. (**B**) Total nucleolar volume per nucleus correlates with NOR copy number in control (top) and reversine-treated (bottom) DLD1 cells. Each black cross represents a nucleus; the red dashed line indicates linear regression; the background color map shows point density (Kernel Density Estimation) (n=42 for control DLD1 cells, n=40 for reversine-treated DLD1 cells). (**C**) Correlation between individual nucleolar volume and number of NORs associated per nucleolus in DLD1 cells. Each point represents an individual nucleolus; Lines represent linear regression; the shaded areas indicate 95% confidence intervals. (n=125 for nucleoli in control DLD1 cells, n=186 for nucleoli in reversine-treated DLD1 cells). (**D-E**) Correlation between individual nucleolar volume and number of NORs associated per nucleolus in SW480 cells (**D**) and HT29 cells (**E**). Each point represents an individual nucleolus; lines represent the linear regression; shaded areas indicate 95% confidence intervals. (n=208 for nucleoli in SW480 cells, n=178 for nucleoli in HT29 cells).

Most NORs remained co-localized with nucleoli (Supplementary Figure 4*C*), consistent with previous findings that NORs on human acrocentric chromosomes are active and associated with nucleoli by default (30). In control cells, the number of NOR signals per nucleus was tightly clustered around 10, consistent with the expected diploid karyotype. By contrast, in reversine-treated cells, we observed a broader distribution of NOR numbers per nucleus, reflecting complex aneuploidy (Supplementary Figure 4*D*). There was a linear correlation between total nucleolar volume per nucleus and the number of NORs per nucleus. In Kernel Density Estimation plots, the number of NORs per nucleus showed a clear rightward shift in the reversine-treated group relative to controls, while total nucleolar volume per nucleus exhibited a corresponding upward shift (Figure 4*B*). This pattern also highlights the increased cell-to-cell heterogeneity in both NOR dosage and nucleolar morphologies following induction of chromosomal instability.

Moreover, there was a strong linear relationship between individual nucleolar volume and the number of associated NORs in both control and reversine-treated cells (Figure 4*C*), further support that the number of NOR associations determines individual nucleolar volumes. We also applied 3D ImmunoFISH to two additional colorectal cancer cell lines which show complex aneuploidy and chromosomal instability at baseline: SW480 and HT29 (31). Similar to reversine-treated DLD1 cells, most NORs were associated with nucleoli by default (Supplementary Figure 5*A-B*; Supplemental Videos 3 and 4) and individual nucleolar volume showed a strong linear relationship with the number of associated NORs (Figure 4*D-E*). This consistent correlation across euploid and heterogeneous aneuploid populations support that NOR dosage mediates nucleolar volume across a spectrum of karyotypes.

### NOR accumulation results in a dominant nucleolus in individual nuclei

Additional analysis of 3D ImmunoFISH data revelaed that reversine-treated DLD1 cells frequently displayed prominent and dominant single nucleoli with numerous co-localized NOR signals per nucleus, whereas control cells more frequently had multiple nucleoli of similar size (Figure 5*A*). In some dominant large-volume nucleoli, the number of NORs concentrated within a single nucleolus exceeded the total NOR count typically present in an entire euploid nucleus (up to 22). These prominent nucleoli are reminiscent of those that can be seen in human malignancies (3).

**Figure 5.**
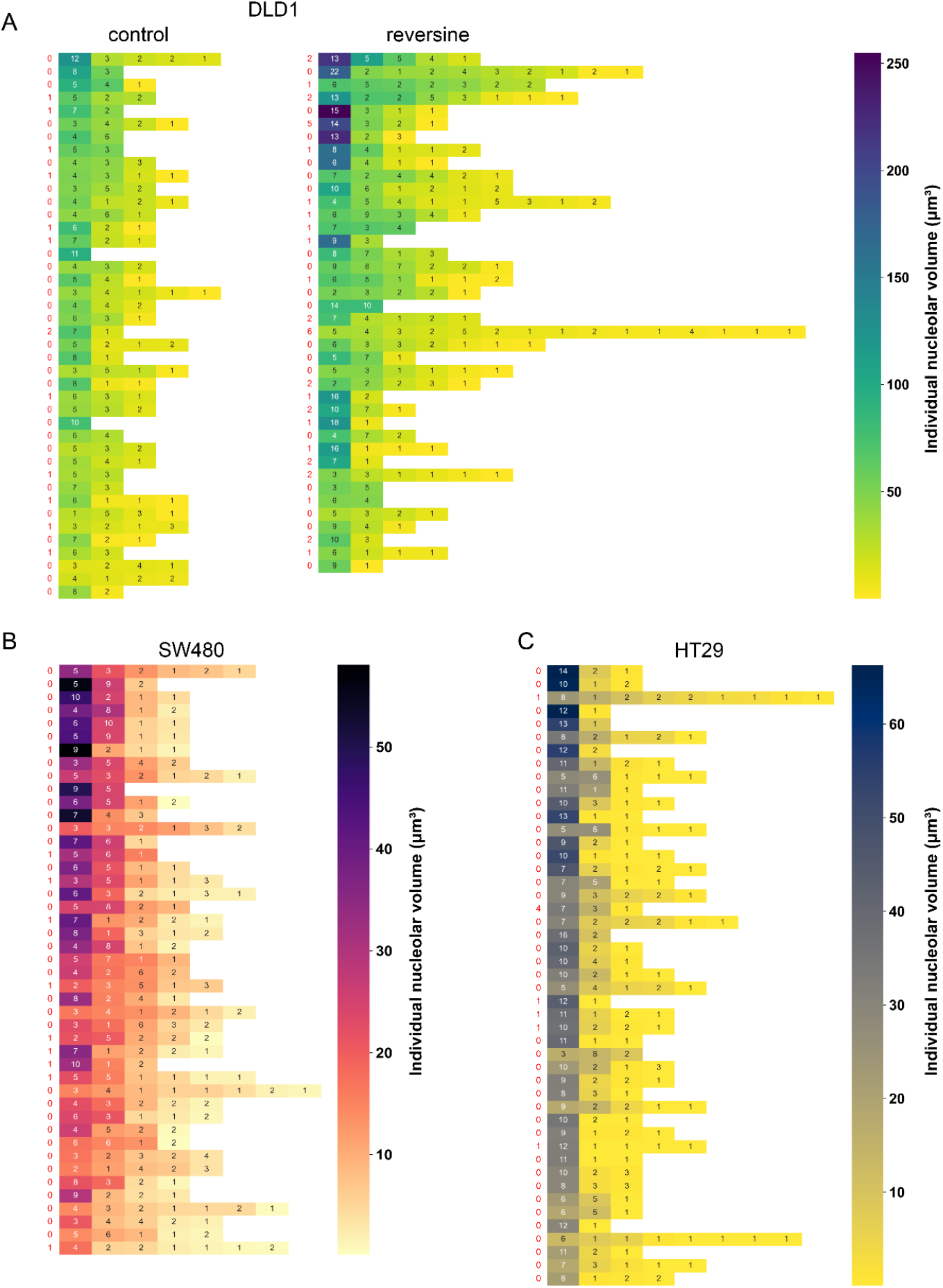
Prominent single nucleoli emerge from NOR accumulation. (**A-C**) Heatmap of individual nucleolar volumes and number of NORs co-localized with individual nucleoli in control and reversine-treated DLD1 cells (**A**), SW480 cells (**B**), and HT29 cells (**C**). Each row represents a single nucleus, with rectangles representing individual nucleoli quantified in the nucleus. Rows are arranged in descending order from highest total nucleolar volume per nucleus (top) to lowest (bottom). The color scale of each rectangle represents the volume of one individual nucleolus, and the number within each rectangle represents the number of 3D co-localized NORs in that nucleolus. The red number to the left of each row represents the number of non-nucleolar associated NORs in the nucleus.

Analysis of SW480 and HT29 cells yielded similar results. When comparing individual nucleoli within the same nucleus, we frequently observed one nucleolus that was markedly larger than the others and harbored the highest number of NOR signals (Figure 5*B-C*). This intra-nuclear pattern was evident in DLD1 controls, DLD1 reversine-treated cells, HT29 cells, and SW480 cells, all of which frequently exhibited dominant nucleoli enriched for multiple NOR signals. Overall, the data support a model in which aneuploidy promotes NOR accumulation and coalescence into dominant nucleoli in cancer.

## Discussion

This study connects aneuploidy to nucleolar remodeling in human cells and provides a new mechanistic explanation for prominent nucleoli frequently seen in cancer. Using MPS1 inhibitor reversine to induce chromosome missegregation, we show that nucleolar number and volume scales with genome content but not with global protein synthesis. Individual nucleolar volumes directly correspond to the number co-localized NOR signals, establishing NOR dosage as a cytogenetic modulator of nucleolar morphologies. Unlike 2D imaging approaches that estimate subcellular dimensions based on projected areas, our 3D imaging pipeline allowed direct quantification of volumetric parameters with high spatial fidelity. These measurements were derived from 3D reconstructions, providing a more biologically accurate assessment of nucleolar morphology and co-localization of NOR signal. Control cells showed a narrow NOR count (near ten) with relative uniform nucleolar volumes, whereas reversine-treated cells produced broader NOR distributions and larger nucleoli as NOR number rose. Prominent single nucleoli containing multiple NORs (sometimes exceeding the total NORs in a euploid nucleus), were common in aneuploid cells, consistent with coalescence of multiple nucleoli.

These results establish NOR dosage as a new contributor to nucleolar morphology and biology, in addition to ribosomal biogenesis, the cellular stress response, and biomolecular condensate dynamics (32) (33) (6). Determining the relative contributions of these factors in various cellular contexts along the benign to malignant spectrum will be important for a more comprehensive understanding of nucleolar size regulation.

These findings have conceptual implications for cancer biology. We observed that the vast majority of NORs are nucleolar-associated across a range of karyotypes, even up to the maximum 40 NORs observed in a single nucleus. This indicates that NOR-nucleolar associations are independent of karyotype, as we did not observe a threshold nucleolar volume above which NORs did not associate with nucleoli. Chromosomal instability and modulation of NOR dosage can have a non-genomic consequence on nuclear condensate architecture, and spatially constrain acrocentric chromosomes at individual nucleoli. Because genome organization is concentrated in and around the nucleolus (34) (35) (36), the formation of dominant nucleoli may alter the local three-dimensional chromatin environment to impact cellular plasticity and increase opportunities for non-random chromosomal rearrangements, perpetuating genomic instability.

Our study has several limitations. Metaphase spread karyotyping was performed on mitotic cells, whereas nucleolar morphologies were measured in randomly chosen interphase cells; thus, the samples may not be identical in chromosomal composition. Moreover, reversine generates a heterogeneous spectrum of aneuploidies, precluding assignments of phenotypes to specific chromosomal gains. These issues could be addressed by combining single-cell karyotype profiling with imaging in the same cells and by using genetic or chemical approaches that bias mis-segregation of specific acrocentric chromosomes or engineered aneuploid cells in our experimental model (37) (38). Finally, our experimental design captured nucleolar phenotypes 24 hours after recovery in reversine-free medium. Future studies will determine the dynamics of NOR-nucleolar associations and nucleolar morphologies across mitotic events.

In summary, these data establish NOR dosage as a cytogenetic mediator of nucleolar number and volume, link aneuploidy and nucleolar biology, and contribute to our understanding of why abnormal nucleolar morphology is a pathognomonic histologic feature of human cancer.

## Materials and Methods

### Cell culture

DLD1 cells were a gift from the laboratory of Dr. Rong Li, and SW480 and HT29 were gifts from the laboratory of Dr. James Eshleman (Johns Hopkins University School of Medicine). DLD1 cells and SW480 cells were maintained in RPMI 1640 medium (Gibco) supplemented with 10% fetal bovine serum (FBS; Sigma), 100 U/mL penicillin, and 100 μg/mL streptomycin (Gibco). HT29 cells were maintained in DMEM (Gibco) with the same supplements. Cells were cultured at 37°C in a humidified incubator with 5% CO_2_.

The human colon organoid line used in this study was derived from freshly resected normal ascending colon tissue from a treatment-naïve colorectal cancer patient, as described previously (21). All procedures were approved by the Johns Hopkins Medicine Institutional Review Board (IRB00125865). Briefly, colon mucosa was separated, finely minced, and incubated in 20 mM EDTA at 37°C on an orbital shaker (200 rpm) for 20 min to release crypts. Crypts were collected by centrifugation and embedded in growth factor-reduced Matrigel (Corning). Colonoids were maintained in 24-well plates (Falcon) or an 8-well chambered coverglass (Nunc LabTek II) in WENR culture medium (ordered from the Johns Hopkins University School of Medicine Single Cell and Transcriptomics Core) as previously described (39), comprised of 50% Wnt3a-conditioned media, 20% R-Spondin1-conditioned media, 10% Noggin-conditioned media and 20% advanced Dulbecco’s modified Eagle’s medium/F12 with 50 ng/mL EGF, supplemented with penicillin, streptomycin, HEPES, GlutaMAX, B27 Supplement, N-acetylcysteine and A83-01. Colonoids were passaged every 6–7 days using TrypLE Express (Gibco) enzymatic dissociation. Matrigel-embedded colonoids were plated at 25 µL per well in 24-well plates or 15 µL per well in 8-well chambers. After polymerization at 37°C for 10 minutes, WENR culture media was added (500 µL for 24-well plates, 400 µL for 8-well chambers). Y-27632 (10 µM, Sigma) was added in the WENR culture media for the first 2 days after passaging to promote cell viability.

### Drug treatment

For both DLD1 cells and colonoids, reversine (Cayman Chemical, Catalogue# 10004412) was added to respective culture media at a final concentration of 500 nM for 24 hours, and an equivalent volume of DMSO (Corning, solvent for reversine) was added in control groups as the vehicle control. For DLD1 cell experiments, reversine was added 2 days after passaging; for colonoid experiments, reversine was added 4 days after passaging. Following treatment, cells and organoids were rinsed three times with phosphate-buffered saline (PBS) to remove residual drug, and fresh culture medium was added for recovery for 24 hours before proceeding to experiments, as previously described (21).

### Metaphase chromosome spreads

DLD1 cells and colonoids were treated with culture medium containing KaryoMAX Colcemid (Gibco, 1:100 dilution) for 8 hours and dissociated using Trypsin (Gibco) or TrypLE Express at 37°C. Single cells were resuspended in 6 mL of prewarmed hypotonic solution (0.56% KCl) and incubated in a 37°C water bath for 10 minutes. Cells were prefixed by adding 1.5 mL of fixative solution (methanol:glacial acetic acid, 3:1) and incubated for 5 minutes at 37°C. After centrifugation at 300 × g for 5 minutes at room temperature, the supernatant was removed, and cells were fixed in 6 mL of fresh fixative for 10 minutes at 37°C. Following another centrifugation, cells were resuspended in 6 mL of fixative and immediately centrifuged without additional incubation.

The cell suspension was dropped onto heat-moisturized glass slides (two drops per slide) and incubated on a heated plate at 75°C for 30 minutes. Slides were then mounted with 20 μL of DAPI-containing mounting medium (Vector Laboratories) and covered with a 22 × 40 mm coverslip. Metaphase spreads were imaged using a Nikon TiE microscope (60 × oil immersion objective).

### Live imaging

DLD1 cells and colonoids were cultured in 8-well chambered coverglass dishes (Nunc LabTek II). For visualization of nuclear and nucleolar structures, samples were stained with DAPI (1 μg/mL) and Nucleolar ID (Enzo, 1:1000 dilution) incubated for 30 minutes at 37°C, then washed with 1x Assay buffer. Live imaging was performed using a Zeiss LSM880 Airyscan laser scanning confocal microscope (40 × water immersion objective). For time-lapse imaging of mitotic DLD1 cells, cells were cultured in the same chambered dishes for 2 days and then imaged by the same microscope for 16 hours. All imaging was conducted at 37°C in a humidified chamber with 5% CO_2_ to maintain physiological conditions.

### Lentivirus

Lentivirus was produced by co-transfecting HEK 293FT cells with the transfer plasmid, packaging plasmid (psPAX2, Addgene #12260), and envelope plasmid (pMD2.G, Addgene #12259). Transfer plasmids used in this study included pLV-H2B-Neon-T2A-mCherry-CAAX-ires-Puro (gift from the laboratory of Dr. Rong Li) (22) and pCA12-pHR-SFFV-BFP-B23 (Addgene #199443).

Virus-containing supernatants were collected at 24 and 48 hours post-transfection, filtered through a 0.45 μm membrane, and concentrated using Lenti-X Concentrator (Takara Bio). Target cells were transduced with concentrated virus in the presence of 8 μg/mL polybrene (Millipore) and selected with puromycin (5 μg/mL) and cell sorter (Sony SH800S).

### OPP

To quantify global protein synthesis, cells were labeled using the Click-&-Go® Plus OPP Protein Synthesis Assay Kit (Vector Laboratories) according to the manufacturer’s instructions. Cells were cultured in an 8-well chambered coverglass (Nunc LabTek II). OPP reagent was diluted 1:1000 in complete culture medium to a final concentration of 20 μM. Cells were incubated with OPP for 30 min at 37°C. Following labeling, cells were fixed with 4% formaldehyde in PBS for 15 min at room temperature, and permeabilized with 0.5% Triton X-100 in PBS for another 15 min. After washing, OPP was labeled by copper-catalyzed click chemistry using AZDye 647 Azide Plus fluorescent dye included in the kit. The reaction cocktail was freshly prepared and added to samples for 20 min at room temperature in the dark. After washing, cells were imaged using Zeiss LSM880 Airyscan laser scanning confocal microscope (40 × water immersion objective). To confirm the specificity of OPP labeling, a subset of cells was pre-treated with 50 μg/mL cycloheximide for 1 hour prior to OPP incubation, which resulted in the loss of OPP signal, verifying inhibition of translation elongation (40).

### 3D ImmunoFISH

DLD1 cells were grown on 3’’ x 1’’ x1 mm slides (VMR VistaVision) in a 10 cm dish and fixed in 4% paraformaldehyde (PFA) for 10 min at room temperature, followed by permeabilization with 0.5% Triton X-100 and 0.5% saponin in PBS for 10 min. Slides were then incubated in 20% glycerol in PBS, snap-frozen in liquid nitrogen, and stored in −80°C.

For DNA FISH, probes targeting the distal junction (DJ) sequences of NORs were amplified from BAC clone CH507-535F5 (BACPAC Genomics, Inc.). Samples were depurinated with 0.1 N HCl for 5 min and equilibrated with 50% formamide in 2 x SSC at 37°C for 15 min. Hybridization mix containing probe was applied to slides, and coverslips were sealed with rubber cement. Slides were denatured at 73°C for 12 min and hybridized at 37°C in a humidified chamber for 48 h. After FISH, nucleoli were stained with an anti-nucleolin antibody (Cell Signaling, 1:1000 dilution), or anti-CDT1 antibody (Cell Signaling, 1:200 dilution) and anti-Geminin antibody (Cell Signaling, 1:800 dilution) for 1 h at room temperature, followed by a secondary antibody conjugated to Alexa Fluor 568 (Invitrogen, 1:1000 dilution) for 1 hour at room temperature (28). Slides were mounted with DAPI-containing mounting medium (Vector Laboratories). Z-stack images were acquired using the Zeiss LSM880 Airyscan laser scanning confocal microscope (40 × oil immersion objective). Volumetric segmentation of nucleoli and quantification of NOR signals per nucleus and per nucleolus were performed using Imaris software (Oxford Instruments, version 10.2.0).

### Quantification and statistical analysis

The number of cells analyzed and the statistical test performed are indicated in each figure legend. All data were collected from two or three independent experiments. Statistical tests and figures were generated using Prism (GraphPad, version 10.6.1) and/or Jupyter Notebook (version 6.5.4). For all statistical tests, a cutoff of p < 0.05 was used to indicate significance.

## Acknowledgments

We would like to acknowledge funding from the Basic Cancer Research Foundation. We would also like to acknowledge Dr. Andrew Ewald for generous access to Imaris software.

## Figures

**Supplementary Figure 1.**
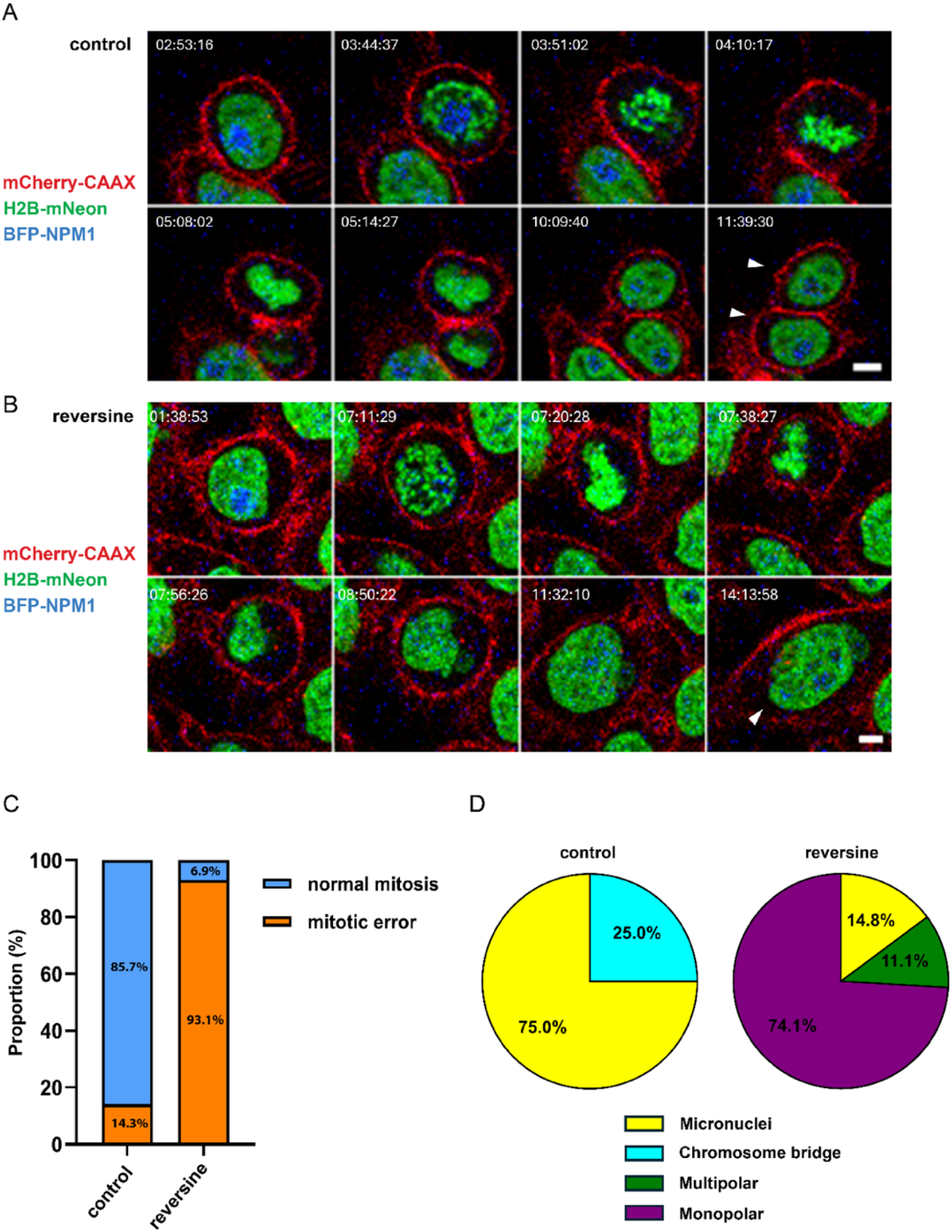
Live imaging of reversine-treated cells shows chromosomal instability and increased mitotic errors. (**A-B**) Representative frames from time-lapse videos of dividing DLD1 cells. Normal mitosis in control DLD1 cells (**A**) and a mitotic error in reversine-treated DLD1 cells (**B**). White arrowheads denote daughter cells. Scale bars, 5 µm. (**C**) Percentage of mitotic errors in dividing DLD1 cells between control and reversine-treated groups. (n=28 for control DLD1 cells, n=29 for reversine-treated DLD1 cells). (**D**) Percentage of types of mitotic errors in dividing DLD1 cells between control and reversine-treated groups. (n=4 for control DLD1 cells, n=27 for reversine-treated DLD1 cells)

**Supplementary Figure 2.**
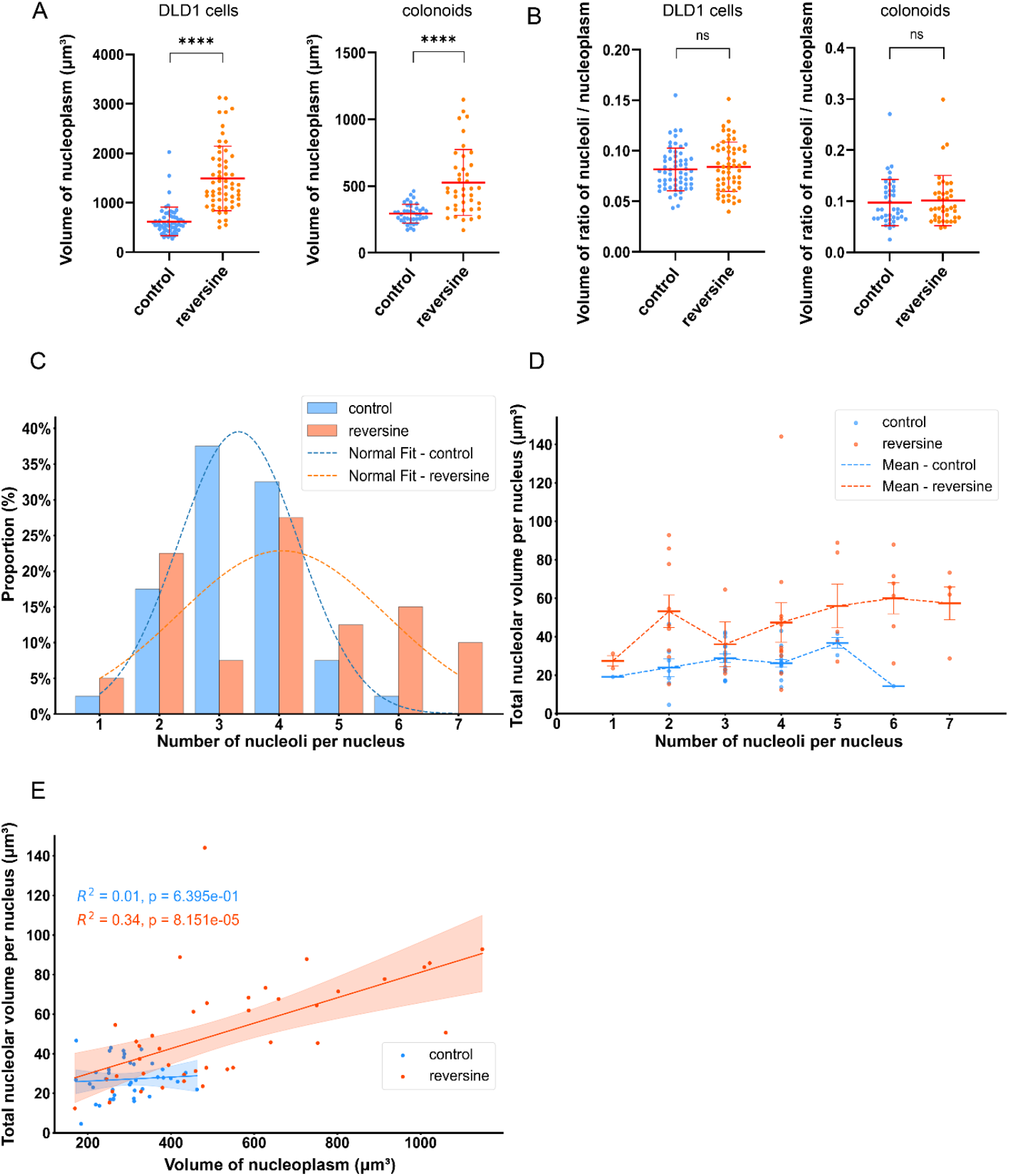
Preserved nucleolar to nucleoplasm ratios and nucleolar morphology changes in reversine-treated lines. (**A**) Volume of nucleoplasm in DLD1 cells (left) and colonoids (right) between control and reversine-treated groups. Lines indicate the mean, and error bars represent the SD. ****p < 0.0001 (Mann-Whitney test). (n=59 for control DLD1 cells, n=58 for reversine-treated DLD1 cells; n=40 for both control and reversine-treated colonoids). (**B**) Ratios of total nucleolar to nucleoplasm volume in DLD1 cells (left) and colonoids (right) between the control and reversine-treated groups. Lines indicate the mean, and error bars represent the SD. ns means not significant (Mann-Whitney test). (**C**) Histogram showing the distribution of the number of nucleoli per nucleus in control versus reversine-treated colonoids. Dashed lines represent the normal distribution fits. (**D**) The total nucleolar volume plotted against nucleolar number per nucleus in colonoids. Each point represents one nucleus. Lines indicate the mean, and error bars represent the SEM. (**E**) Linear correlation analysis showing the total nucleolar volume versus nucleoplasm volume per nucleus in colonoids. Each point represents an individual nucleus; the lines represent linear regression; shaded areas indicate 95% confidence intervals.

**Supplementary Figure 3.**
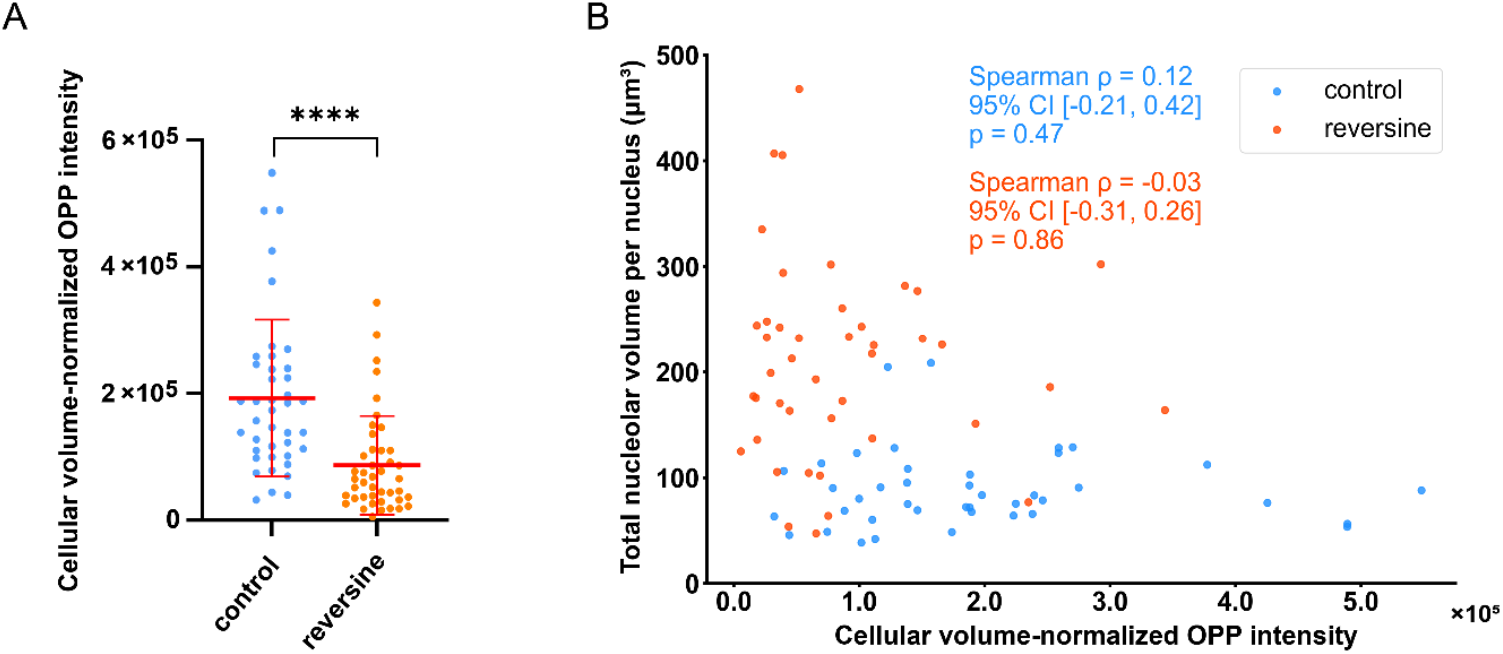
OPP fluorescence intensity normalized to cellular volume. (**A**) Quantification of cellular volume–normalized OPP fluorescence intensity in control and reversine-treated DLD1 cells. Lines indicate the mean, and error bars represent the SD. ****p < 0.0001 (Mann-Whitney test). (n=41 for control DLD1 cells, n=43 for reversine-treated DLD1 cells). (**B**) Scatter plots showing the relationship between cellular volume–normalized OPP intensity and total nucleolar volume per nucleus. Each point represents an individual cell. Spearman rank correlation coefficients (ρ), two-sided p values, and 95% bootstrap confidence intervals are indicated.

**Supplementary Figure 4.**
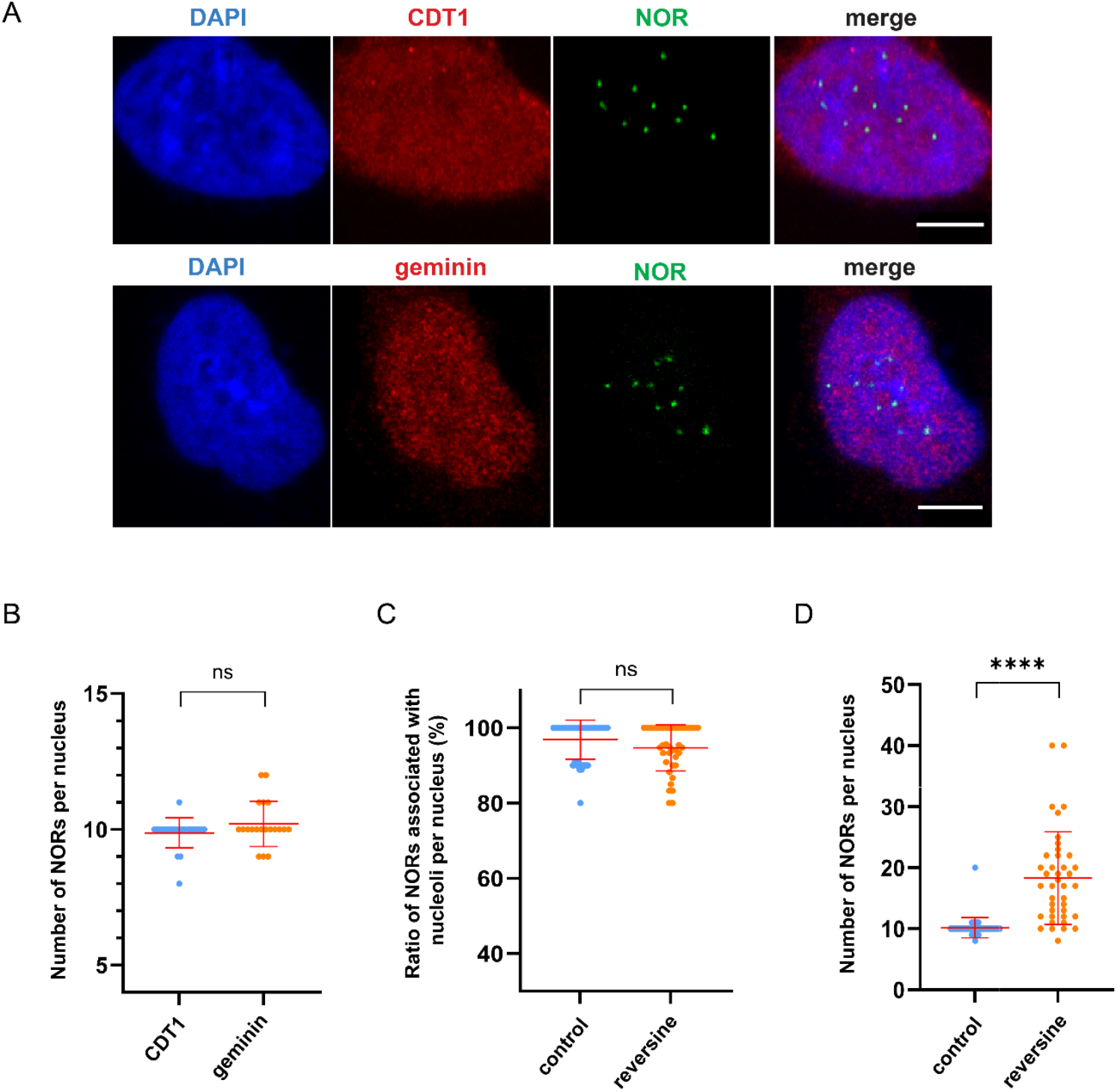
3D ImmunoFISH of DLD1 cells. (**A**) Representative images of DLD1 cells labeled with distal junction (DJ)-specific FISH probes targeting NOR loci and anti-CDT1 antibody (top) and anti-Geminin antibody (bottom). Scale bars, 5 µm. CDT1 is a marker for the G1 phase of the cell cycle, while geminin is a marker for the S/G2 phases. (**B**) Quantification of the number of NOR foci per nucleus between CDT1 positive and Geminin positive DLD1 cells. Lines indicate the mean, and error bars represent the SD. ns means not significant (Mann-Whitney test). (n=23 for CDT1-positive cells, n=20 for geminin-positive cells).(**C**) Percentage of NORs associated with nucleoli per nucleus between control and reversine-treated DLD1 cells. Lines indicate the mean, and error bars represent the SD. (n=42 for control DLD1 cells, n=40 for reversine-treated DLD1 cells). (**D**) Quantification of number of NORs per nucleus between control and reversine-treated DLD1 cells. Lines indicate the mean, and error bars represent the SD. ****p < 0.0001 (Mann-Whitney test) (n=42 for control DLD1 cells, n=40 for reversine-treated DLD1 cells).

**Supplementary Figure 5.**
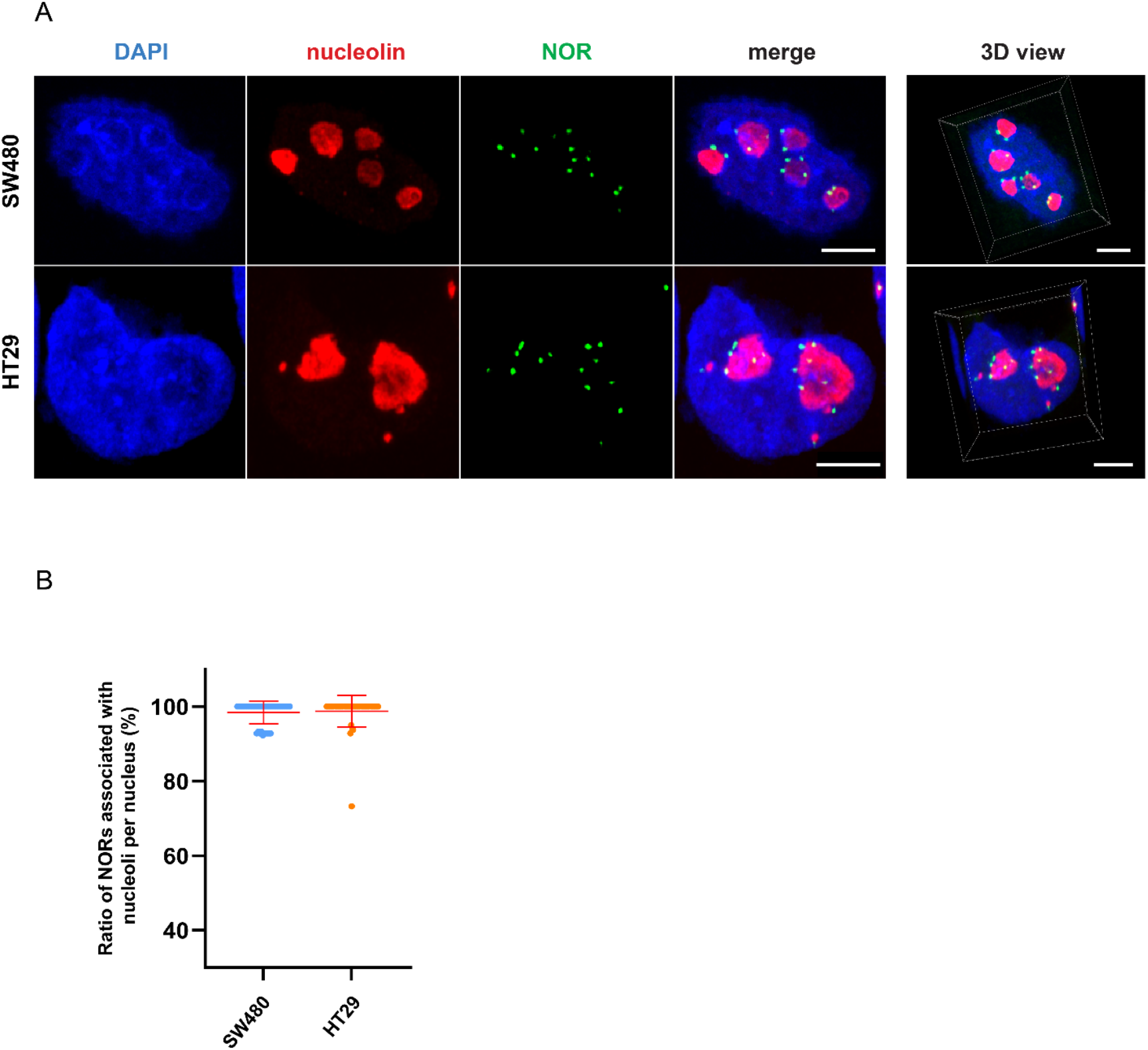
3D ImmunoFISH of SW480 and HT29 cells. (**A**) Representative 3D ImmunoFISH images of SW480 and HT29 cells labeled with anti-nucleolin antibody (red, nucleoli) and distal junction (DJ)-specific FISH probes targeting NOR loci (green). DNA is stained with DAPI (blue). Scale bars, 5 µm. (**B**) Percentage of NORs associated with nucleoli per nucleus in SW480 cells and HT29 cells. Lines indicate the mean, and error bars represent the SD. (n=45 for SW480 cells, n=47 for HT29 cells).

